# Glucose signaling is important for nutrient adaptation during differentiation of pleomorphic African trypanosomes

**DOI:** 10.1101/346601

**Authors:** Yijian Qiu, Jillian E. Milanes, Jessica A. Jones, Rooksana E. Noorai, Vijay Shankar, James C. Morris

**Author notes:** Corresponding Author: (JCM).

## Abstract

The African trypanosome has evolved mechanisms to adapt to changes in nutrient availability that occur during its lifecycle. During transition from mammalian blood to insect vector gut, parasites experience a rapid reduction in environmental glucose. Here we describe how pleomorphic parasites respond to glucose depletion with a focus on parasite changes in energy metabolism and growth. Long slender bloodstream form parasites are rapidly killed as glucose concentrations fall, while the short stumpy bloodstream form parasites persist to differentiate into the insect stage procyclic form parasite. The rate of differentiation was slower than that triggered by other cues but reached physiological rates when combined with cold shock. Both differentiation and growth of resulting procyclic form parasites were inhibited by glucose and its non-metabolizable analogs in a concentration dependent manner. Procyclic form parasites differentiated from short stumpy form parasites in glucose depleted medium significantly upregulated gene expression of amino acid metabolic pathway components when compared to procyclic forms generated by cis-aconitate treatment. Additionally, growth of these parasite was inhibited by the presence of either glucose or 6-deoxyglucose. In summary, glucose transitions from the primary metabolite of the blood stage infection to a negative regulator of cell development and growth in the insect vector, suggesting that the hexose is not only a key metabolic agent but is also an important signaling molecule.

**Author Summary:** As the African trypanosome, *Trypanosoma brucei*, completes its lifecycle, it encounters many different environments. Adaptation to these environments includes modulation of metabolic pathways to parallel the availability of nutrients. Here, we describe how the blood-dwelling lifecycle stages of the African trypanosome, which consume glucose to meet their nutritional needs, respond differently to culture in the near absence of glucose. The proliferative long slender parasites rapidly die, while the non-dividing short stumpy remains viable and undergoes differentiation to the next lifecycle stage, the procyclic form parasite. Interestingly a sugar analog that cannot be used as an energy source inhibited the process. Furthermore, the growth of procyclic form parasite that resulted from the event was inhibited by glucose, a behavior that is similar to that of parasites isolated from tsetse flies. Our findings suggest that glucose sensing serves as an important modulator of nutrient adaptation in the parasite.

## Introduction

Organisms that occupy multiple biological niches must adapt to different environments. Such is the case for the vector-borne African trypanosome, *Trypanosoma brucei*, a kinetoplastid parasite that is the causative agent of African sleeping sickness. This parasite, which is transmitted by tsetse flies, undergoes a series of developmental steps that yield lifecycle stages which are uniquely adapted for life in the distinct hosts. These adaptations include alterations to metabolic pathways that parallel differences in substrate availability and expression of distinct surface molecules that are required for successful colonization of the new environment.

In *T. brucei* development, differentiation events occur in both the mammalian host and insect vector. As their density increases in the vertebrate bloodstream, long slender (LS) blood form parasites perceive a quorum-dependent parasite-derived signal that triggers differentiation into short stumpy (SS) blood form parasites, a non-dividing form arrested in G0 of the cell cycle (1). When these SS parasites, which are pre-adapted for life in the tsetse fly vector (2), are engulfed by a tsetse fly during a blood meal, they quickly differentiate into dividing procyclic form (PF) parasites that are competent for completion of the lifecycle in the fly.

Development is carefully coordinated with environmental setting, ensuring that the appropriate lifecycle stage is initiated in the correct host and tissue. The ability to perceive and respond to the environment requires detection of cues that trigger signaling pathways to modulate gene expression. SS trypanosomes are exposed to an array of potential signals including fluctuating temperatures, exposure to digestive processes in the fly gut, and interaction with other trypanosomes. Additionally, there are an assortment of small molecules generated while the blood meal vehicle in which the SS parasites reside is digested by the fly.

In resolving these sensing pathways, potential cues associated with development have been tested. Exposure of SS parasites to a cold shock, specifically a change in environmental temperature of more than 15°C, has been shown to trigger a nearly immediate and reversible expression of the PF surface molecule EP procyclin (3). However, these cells failed to grow. As cell growth is critical feature of the transition from SS to PF, this observation suggested that that cold shock alone was insufficient for complete initiation of the developmental program. Notably, cold shock also triggered the expression of a family of carboxylate transporter proteins called “proteins associated with differentiation (PAD)” that have been implicated in the differentiation response triggered by a distinct cue, citrate and cis-aconitate (CCA). Without exposure to cold shock, a high concentration of CCA (6 mM) is required to initiate differentiation. However, exposure of SS parasites to cold shock, which results in increased surface expression of PAD2, triggers EP procyclin expression at extremely low levels of cis-aconitate (0.6-6 µM) (3,4). The tsetse fly midgut contains similar levels of citrate (15.9 µM) and this carboxylic acid has been found to mirror the impact of CCA in SS to PF development, an observation that defines CCA as a potentially physiologically relevant cue (3,5).

Additional cues may be associated with cold shock and citrate that enhance differentiation from SS to PF parasites. These include exposure to mild acid or protease treatment (5-9). Treatment of SS parasites with either of these cues rapidly initiated EP procyclin expression (∼2 hours), but the response mechanisms were different. Mild acid treatment, much like exposure to high levels of CCA, led to phosphorylation of TbPIP39, a phosphatase component of the CCA differentiation cascade (10). This phosphorylation event indicates activation of this differentiation pathway, which was not observed after protease treatment. This suggested the response was due to a distinct signaling pathway (9).

One aspect of differentiation is the adaptation to nutrients available in a particular host. In mammalian blood, LS parasites are exposed to ∼5 mM glucose and the hexose serves as a critical carbon source for this form of the parasite. When bloodstream form parasites are taken up by a feeding tsetse fly they experience a rapid drop in glucose concentration, with the sugar depleted from the blood meal in ∼15 minutes (11). In this environment, SS parasites persist and differentiate into PF parasites. Reflecting the reduced glucose levels in the environment, PF parasites complete the activation of metabolic pathways that were initiated in the SS lifecycle stage required for metabolism of amino acids like proline and threonine (12-14).

Glucose is a critical carbon source for the African trypanosome but its role in development is unresolved. Manipulation of glucose levels *in vitro* has been shown to alter developmental patterns and gene expression, suggesting a role for the sensing of the sugar in parasite development. Cultured monomorphic bloodstream form (BF) parasites that do not differentiate into SS could be prompted to differentiate into PF parasites by removal of glucose and addition of glycerol to the growth medium (15). The low survival and differentiation rates, along with the phenomena being noted in a developmentally defective strain, however, call into the question the biological relevance of glucose as a single cue to initiate this process. Additional evidence that glucose sensing plays a potential role in development includes the observation that partial inhibition of glycolysis in monomorphic BF with phloretin, a plant-derived dihydrochalcone, or 2-deoxy-glucose (2-DOG), a glycolytic poison, triggered genomewide transcriptome changes that resembled expression patterns found in SS to PF parasite differentiation. These transcriptome changes included the upregulation of EP procyclin and many genes involved in energy metabolism (16,17). Last, PF parasites were found to alter surface molecule expression in a glucose-dependent manner (18,19) in a process that is regulated by mitochondrial enzymes (19). These studies suggest that while glucose manipulation is not sufficient for initiation of a rapid differentiation program in BF parasites, it may play a role in development across several lifecycle stages.

Here, we present evidence that culturing SS parasite in medium with minimal glucose (∼5 μM) triggers differentiation to PF parasites. The rate of this glucose-responsive differentiation as measured by EP procyclin expression and initiation of growth was slower than rates observed with other known cues of differentiation. However, the rate was enhanced to physiologically relevant levels when cells were also exposed to a brief cold shock, which is a known potentiator of other differentiation cues. The resulting PF parasites upregulated genes involved in amino acid metabolism to a greater extent than those differentiated through CCA treatment, indicating a potential role for glucose in nutrient adaptation.

## Results

### Bloodstream form parasite rapidly deplete glucose from the environment

Responses to new environments are particularly important to parasitic microbes that inhabit different hosts during their lifecycles. As African trypanosomes transition from the glucose-rich blood of the mammalian host to the tsetse fly gut, they undergo a marked change in environment. One major change is the rapid (∼15 min) depletion of environmental glucose, the primary carbon source used to generate ATP during bloodstream infection (11). This reduction in available glucose is at least in part due to the metabolic activity of the parasites in the blood meal. Both pleomorphic LS and SS parasites isolated from infected rodents rapidly consume glucose *in vitro* with 0.5 mM glucose being nearly depleted from culture medium after a single day by both lifecycle stages (Fig. 1A). After one day, glucose levels were reduced to 37 ± 0.70 and 62 ± 0.60 µM for LS and SS, respectively and the hexose concentration continued to fall on day two, reaching 1.7 ± 0.70 and 15 ± 0.90 µM. The precipitous decline in glucose availability had an impact on LS parasite viability even though the potential carbon sources proline and threonine were included in the medium. While more than 80% of the LS parasites were dead after two days of culture under the very low glucose conditions, SS parasites were less sensitive with >80% of the population viable after the same period (Fig. 1B).

**Fig 1.**
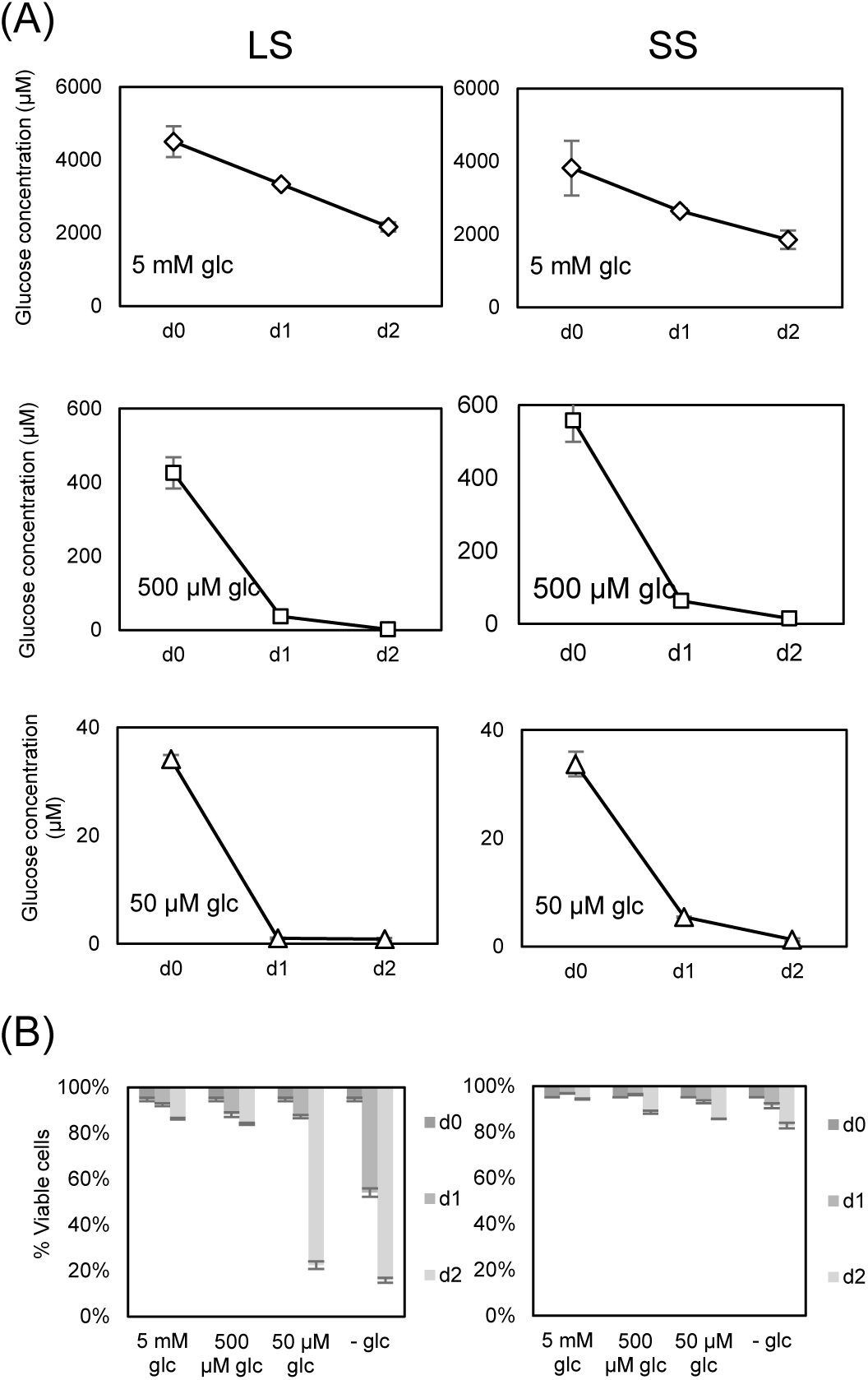
Slender form (LS) and short stumpy form (SS) *T. brucei* rapidly deplete glucose. Parasites isolated from rodent buffy coats by chromatography were washed extensively in PBS and then resuspended (4 × 10^5^/cells/mL for LS and 5 × 10^5^ cells/mL for SS) in RPMIθ supplemented with proline and threonine and different concentrations of glucose. LS parasites (left column) were isolated after four days of infection and made up nearly 100% of the parasite population as determined by microscopy, while the SS samples (right columns) contained a mixture of SS (∼90%) and LS (∼10%) parasites as scored by cytometry of PAD1-labeled parasites (not shown). (A) Glucose concentrations in the medium were measured through time as described in the Materials and Methods and standard deviation from experiments performed in triplicate indicated. (B) Parasite viability was scored by propidium iodide staining.

### Impact of glucose depletion on the differentiation and proliferation of pleomorphic SS and LS and monomorphic BF parasites

Monomorphic BF parasites cultured in very low glucose medium supplemented with glycerol undergo an inefficient differentiation to PF parasites (15). This process is likely to be a response of monomorphic BF parasites to metabolic stress instead of an orchestrated program in response to a developmental cue. To determine if glucose depletion had a similar impact on cell development in different mammalian lifecycle stages, pleomorphic LS and SS form parasites were incubated in SDM79θ, a very low glucose (∼5 µM) PF culture medium that contains amino acids but that lacks glycerol. Culture of pleomorphic SS parasites in SDM79θ under conditions that would normally support PF *in vitro* growth (27°C, 5% CO_2_) led to detectable parasite outgrowth by day 3 (Fig. 2A, open circles). This rate of outgrowth was considerably slower than that generated by known differentiation cues, which we have termed glucose-responsive slow differentiation (GRS differentiation). SS parasites maintained in SDM79θ supplemented with 5 mM glucose (filled circles) did not appreciably grow. While the highly motile and proliferative LS pleomorphic parasites are superficially most similar to the BF monomorphic cells, the LS parasites were different in that they did not tolerate culture in glucose depleted conditions even when glycerol was added to the medium (S1 Fig).

**Fig 2.**
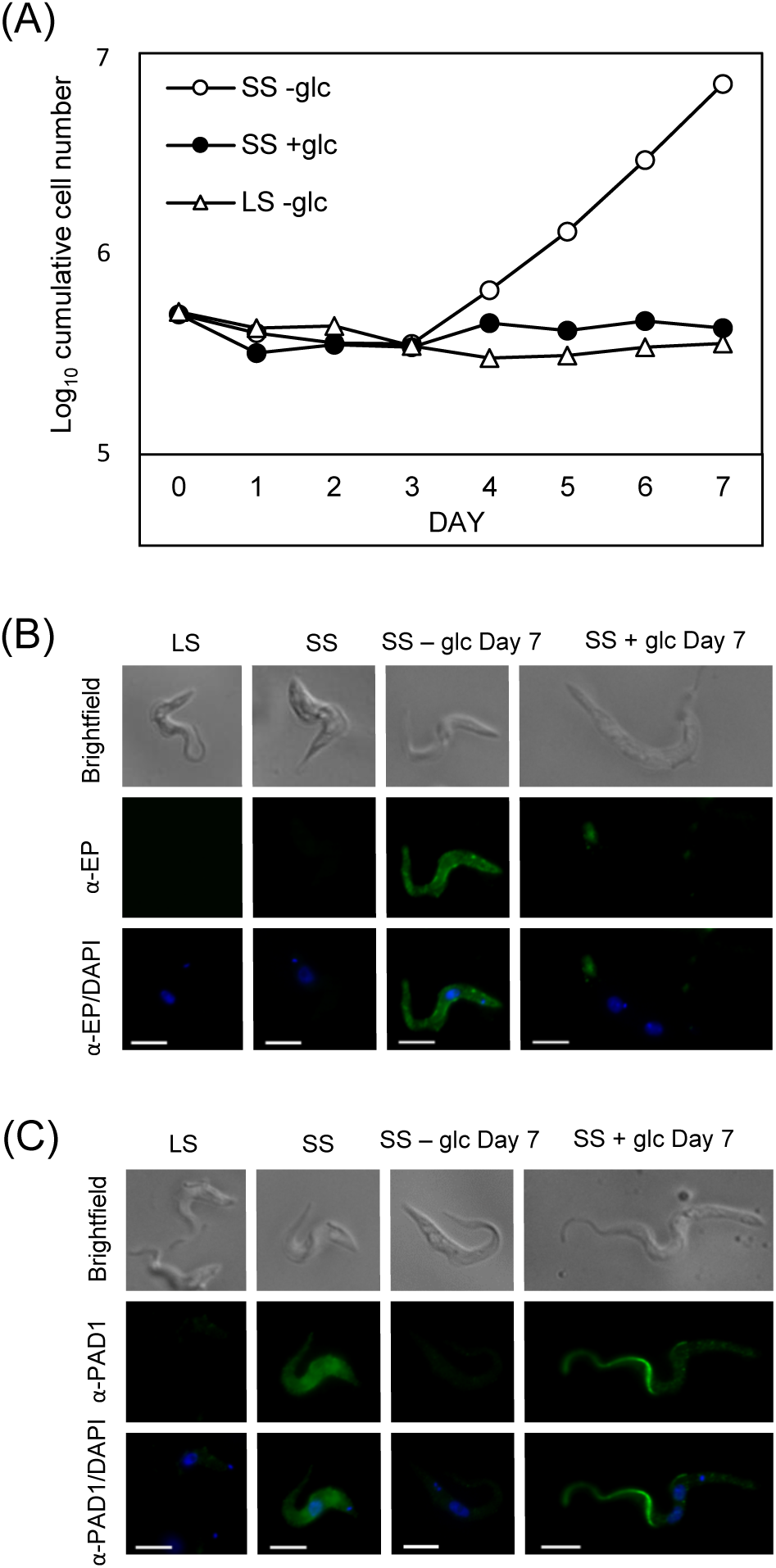
*T. brucei* SS growth and surface molecule expression are influenced by environmental glucose availability. (A) LS and SS parasites isolated from rodents were washed extensively and resuspended in very low glucose PF medium, SDM79θ (-glc, ∼5 μM), supplemented with (5 mM) or without additional glucose at 27°C and growth monitored. Growth curves are representative of at least two assays. (B and C) IF analysis of parasites after incubation for seven days with or without glucose. Fixed parasites were visualized by epifluorescence microscopy using antiserum against EP procyclin (B) or PAD1 (C). DAPI was added with anti-fade reagent to stain the nucleus and kinetoplast DNA in all samples. Scale bar = 5 μm.

SS forms in the mammalian bloodstream are arrested in the cell cycle and do not divide, suggesting that the cell proliferation observed after incubation in SDM79θ was a consequence of these SS differentiating to PF. To confirm this, parasites were probed with antiserum specific to the PF marker protein, EP procyclin, and analyzed by immunofluorescence (IF) (Fig. 2B). IF analysis of growing cells revealed that the parasites were morphologically similar to PF cells, expressing a surface coat of EP procyclin not observed in SS or in SS cultured in the presence of glucose (Fig. 2B).

As an additional marker for development, parasites were probed with antiserum specific for PAD1, a marker for SS parasites, and the highest signal was observed on the surfaces of SS parasites freshly isolated from rodent blood, as has been described (Fig. 2C) (4). Growing parasites generated by culture without glucose led to reduced PAD1 labeling, while the majority of those cultured in the presence of glucose for the same period died. However, the few that persisted frequently harbored multiple nuclei, accumulated PAD1 signal in the flagellum, and became elongated (Fig. 2C). The inability of these cells to progress further along the lifecycle suggests that glucose is a negative regulator of cellular development in the SS parasite. The outgrowth of the normally quiescent SS parasites together with EP procyclin expression and reduction of PAD1 labeling was only observed in the absence of glucose. The majority of the cells remained viable throughout the treatment and the resulting parasites proliferated rapidly as long as they were maintained in very low glucose medium. Together, the expression of PF parasite markers and resumption of growth was indicative of differentiation to the insect form lifecycle stage (S4 Fig).

### GRS differentiation rates reach physiologically relevant rates when combined with cold shock

Triggers that initiate SS to PF lifecycle stage differentiation have been widely studied (3-9). While differentiation of SS parasites as a result of glucose depletion alone was delayed when compared to other described triggers, the parasite is likely exposed to multiple environmental conditions in the tsetse fly that could influence this rate. To investigate the effect of additional cues on GRS differentiation, rates of surface molecule expression and growth were monitored (Fig. 3). Exposure of SS parasites to cold shock in combination with citrate levels found in the tsetse fly midgut (15.9 μM, (5)) triggered EP procyclin expression after three hours and cell growth after four days (Fig. 3A and B). The doubling time of these cells between days 4 and 7 was ∼64.5 hours (Fig. 3C, asterisks). Similarly, the combination of cold shock and the near-absence of glucose yielded EP procyclin expression after three hours and cell proliferation was noted within one day. In addition to proliferation, these cells also grew more rapidly compared to those where citrate was added, with a doubling time of ∼21.5 hours (open triangles) between days 4 and 7.

**Fig 3.**
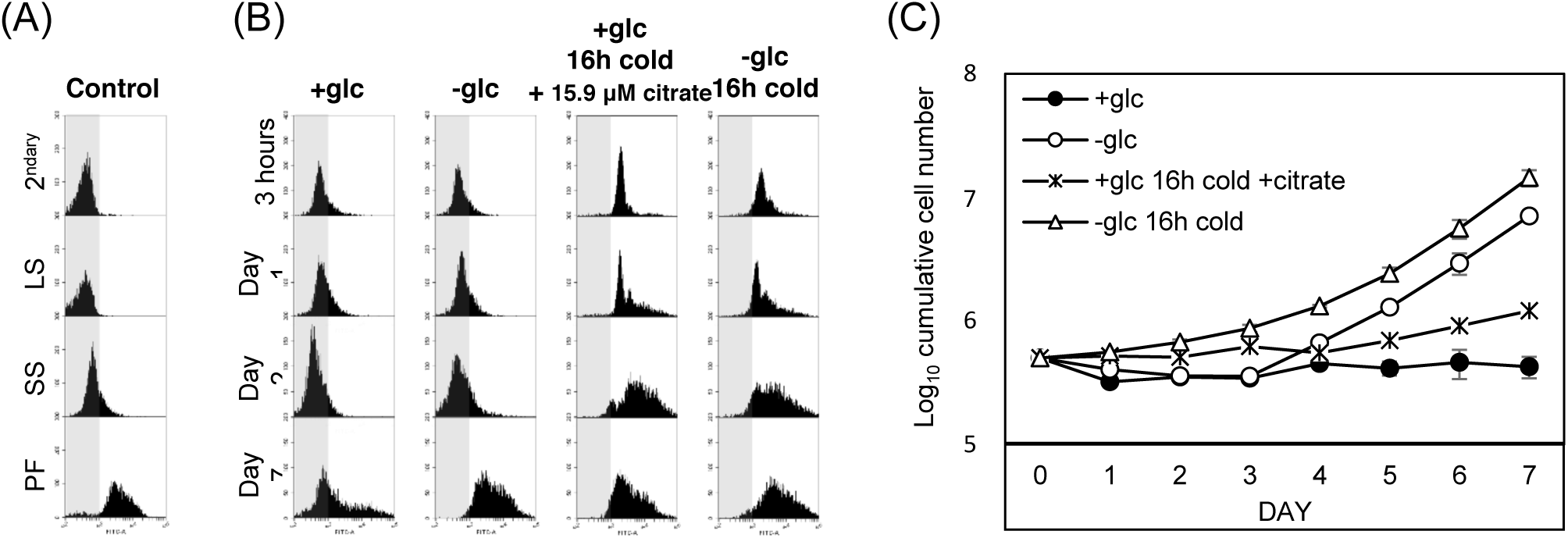
The near-absence of glucose is synergistic with cold treatment in triggering differentiation from SS to PF. Flow cytometry of (A) control experiments (secondary alone (2^ndary^); LS, SS, and PF stained with antiserum to EP procyclin) and (B) SS parasites after culture at 37°C or exposure to 20°C for 16 hours in HMI9 prior to washing and resuspension in SDM79θ supplemented with 5 mM glucose (+glc) or with physiological levels of citrate (15.9 µM). Cells were fixed at the indicated times, stained with antiserum to EP procyclin, and scored by cytometry (5,000 cells/assay) (C) Growth of parasites exposed to cold shock (20°C) or 37°C for 16 hours prior to initiation of this experiment with combinations of other environmentally-relevant cues (at 27°C). Bars indicate standard deviation in triplicate assays.

### Mitochondrial metabolism is required during GRS differentiation

SS parasites can metabolize the abundant glucose in the blood of the mammalian host much like LS parasites. However, SS parasites also express genes that prepare them for life in the glucose-poor environment of the fly gut by upregulating genes required for mitochondrial metabolic functions, including cytochrome C oxidase subunits (20) and respiratory chain complex I genes (21). To test the potential contribution of mitochondrial amino acid metabolism to differentiation, parasite growth was scored after manipulating the available amino acids in the very low glucose (∼ 5 µM) bloodstream form medium RPMIθ. When SS parasites were cultured in medium that included proline and threonine for two days prior to being transferred into glucose-free PF medium that had abundant amino acids (SDM79θ), most cells remained viable and differentiation occurred (Fig. 4A, open circles). Inclusion of threonine alone did not support parasite differentiation (open triangles) while culture in proline alone (open squares) led to a slightly delayed entry into growth when compared to medium with both proline and threonine. RPMIθ medium lacking glucose and both amino acids did not support parasite viability (x symbol), with a greater than 80% loss of SS parasite viability on day one (Fig. 4B). Inhibition of oxidative phosphorylation with the antibiotic oligomycin, an ATP synthase inhibitor, prevented differentiation and was toxic to SS parasites cultured in very low glucose (Fig. 4C, open squares and Fig. 4D). These observations indicated that amino acid metabolism is likely important for satisfying the metabolic needs of the trypanosomes in a minimal glucose environment during GRS differentiation. The ability to adapt to the new carbon source in the absence of glucose is limited to SS parasites, as LS forms are not viable under the same conditions.

**Fig 4.**
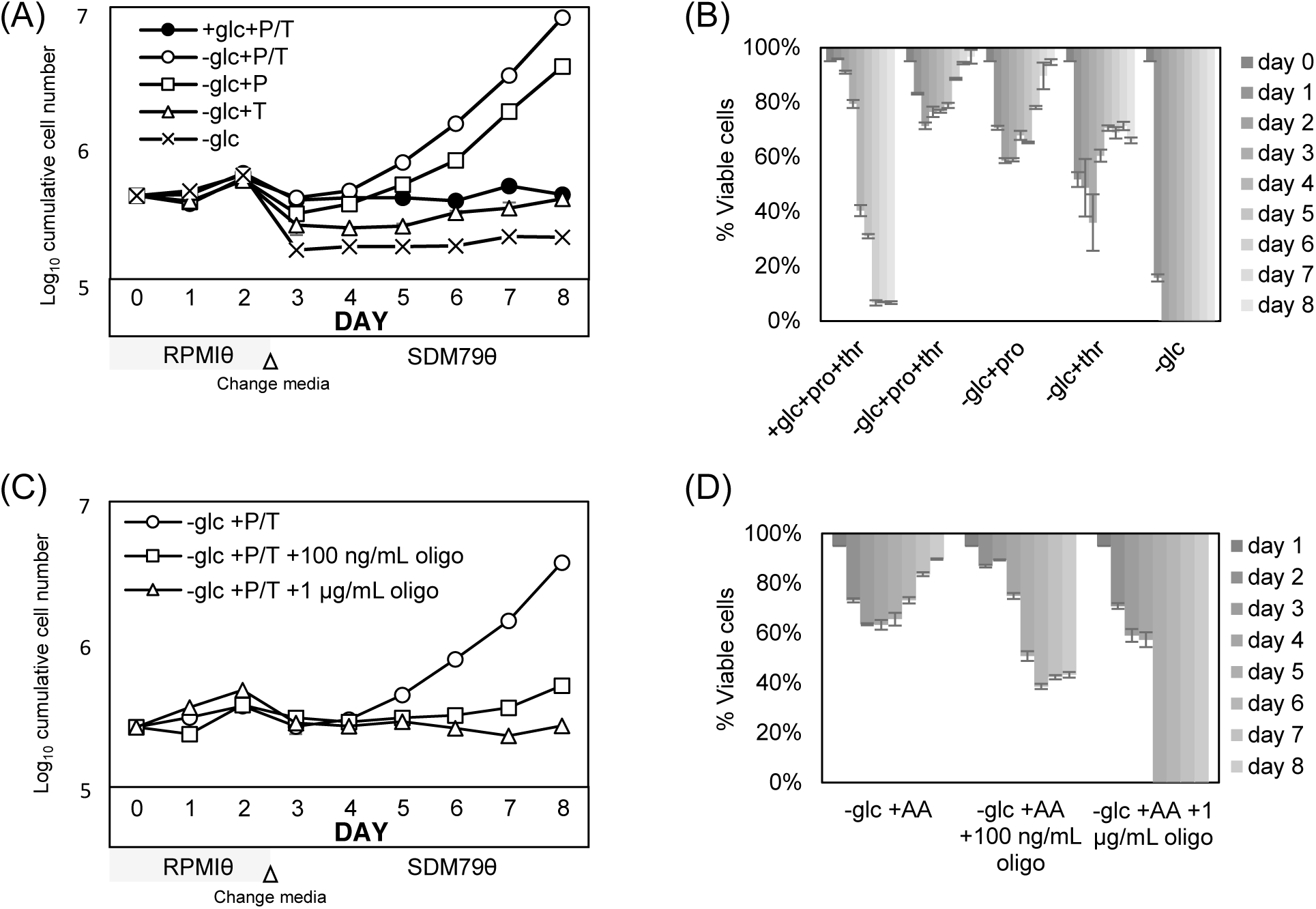
Amino acids are required for completion of SS differentiation and cell viability in very low glucose medium. SS parasites cultured in bloodstream form very low glucose medium, RPMIθ (which allowed manipulation of amino acid content) supplemented with or without glucose, proline (P, 4.6 mM), threonine (T, 3.4 mM), or proline and threonine for two days followed by transfer to amino acid-replete SDM79θ were scored for (A) outgrowth and (B) viability by flow cytometry and propidium idodide (PI) staining. (C) Assessment of parasite outgrowth and (D) viability after oxidative phosphorylation inhibition by treatment with oligomycin (either 100 ng/mL or 1 µg/mL) for two days in RPMIθ with proline and threonine (AA) followed by transfer to SDM79θ to provide amino acids for PF parasites. Bars indicate standard deviation in triplicate assays.

### Glucose inhibits GRS differentiation through a mechanism distinct from glycolysis

In *Saccharomyces cerevisiae*, nutrient adaptation responses to glucose availability are mediated through non-metabolic pathways that transduce extracellular or intracellular glucose levels into recognizable signals (22). However, the importance of glucose to BF parasite metabolism raises the possibility that culture in very low concentrations of the carbon source could initiate a stress response related to either insufficient ATP production by glycolysis or a failure to synthesize required glycolytic intermediates.

To elucidate the role of glycolysis on the GRS differentiation response in *T. brucei*, parasites were cultured with a variety of sugars at different concentrations and their impact on differentiation was scored. First, conditions were tested to determine if they met the metabolic needs of the glycolysis-dependent LS parasites. As anticipated, 2-DOG, which is phosphorylated by hexokinase and then inhibits downstream glycolysis, was toxic to LS parasites. Similarly, 6-deoxy-glucose (6-DOG), an analog of glucose that cannot be phosphorylated by hexokinase, and the five-carbon sugar xylose did not support LS viability (S2 Fig).

SS parasite outgrowth paralleled the depletion of glucose from the medium, although the minimum concentration of glucose required to trigger this response is unclear. When SS parasites were seeded into RPMIθ supplemented with increasing concentrations of glucose (0, 5, and 50 µM), outgrowth was noted at about the same time regardless of the concentration of glucose (S3 Fig). Confounding this experiment, however, SS parasites rapidly metabolize glucose to low levels in culture (Fig. 1), making the assessment of the critical glucose concentration needed to initiate outgrowth difficult.

Non-metabolizable glucose analogs were considered next, in part because SS parasites were anticipated to be less sensitive than LS to these compounds given their ability to persist in the near-absence of glucose (Fig. 1). Incubation with 2-DOG, much like glucose, prevented SS GRS differentiation (Fig. 5A), but this compound was acutely toxic to the SS parasites with >70% lethality after two days of exposure (Fig. 5B). The compound 6-DOG is not a substrate for cellular hexokinases and thus cannot be phosphorylated to enter glycolysis. Similar to 2-DOG and glucose, culturing SS parasites with 5 mM 6-DOG prevented differentiation (Fig. 5A). While incubation in 50 µM 6-DOG prevented outgrowth, 5 µM compound had no impact on cells (Fig. 5C). High concentrations of both glucose and 6-DOG were gradually toxic to SS parasites, with near-complete loss of viability after four days in 5 mM of either compound (Fig. 5D). Lastly, xylose had no inhibitory impact on SS parasite GRS differentiation (Fig. 5A) and, like culturing in very low glucose, was minimally toxic to the SS parasites (Fig. 5B).

**Fig 5.**
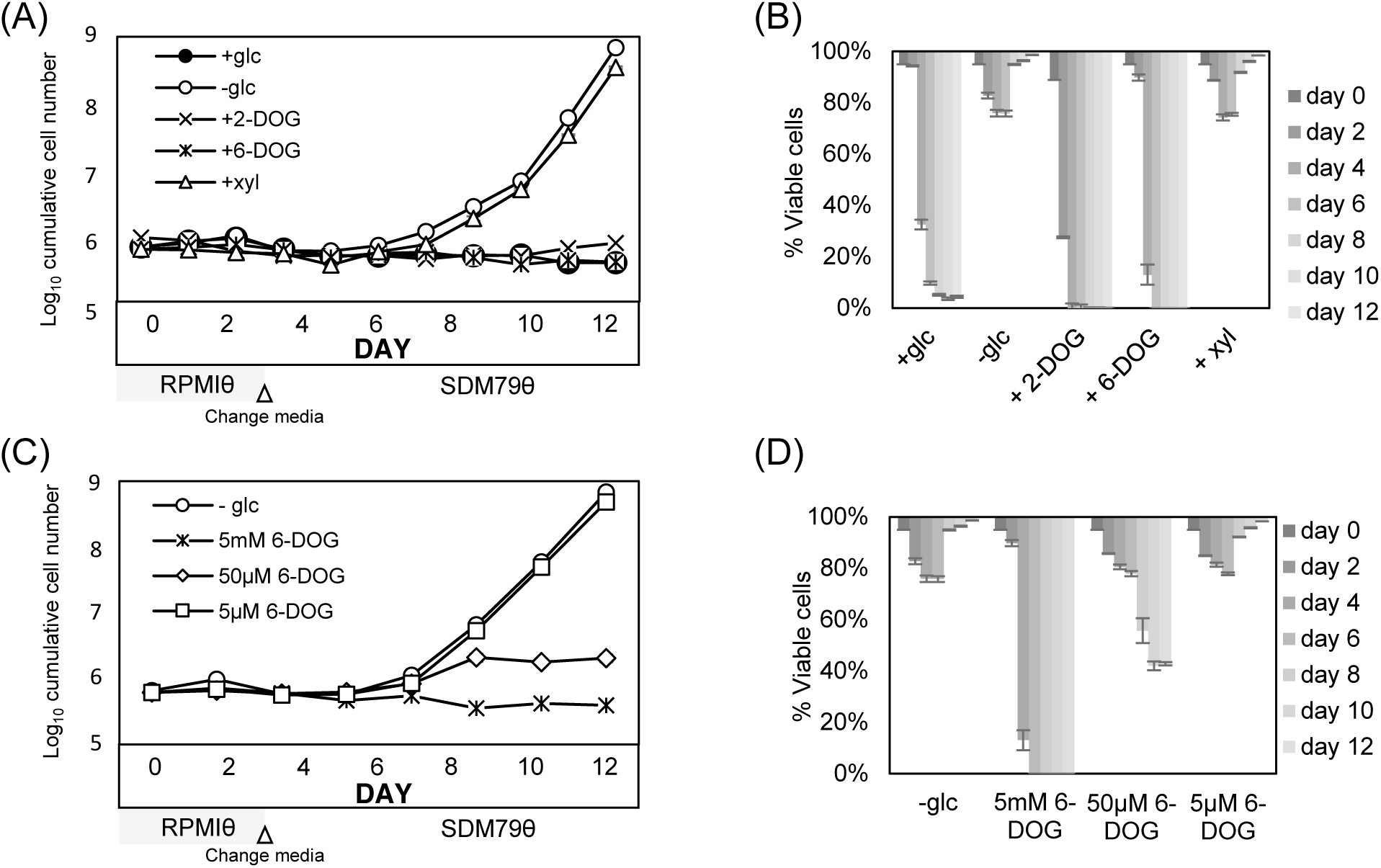
Glucose inhibition of differentiation is independent of glycolysis. (A) Outgrowth and (B) cellular viability of SS cultures after treatment with 5 mM of the indicated compound s (xyl, xylose; 2-DOG, 2-deoxyglucose; 6-DOG, 6-deoxyglucose). SS parasites were washed and resuspended (5 × 10^5^/mL) in RPMIθ with proline and threonine for the first two days and then transferred to the very low glucose PF medium SDM79θ supplemented with 5 mM of the indicated compound and growth monitored. (The transfer was necessary to provide amino acids required for PF viability.) (C) Impact of varied levels of 6-DOG on SS outgrowth and (D) viability by flow cytometry and propidium iodide (PI) staining. Concentrations of 6-DOG were maintained throughout the experiment. Bars indicate standard deviation in triplicate assays.

### Pre-adaption of SS cells at the transcriptome level

SS parasites occupy two extremely different niches (the glucose-rich mammalian blood and the glucose-poor fly midgut) and they express metabolic characters that would allow survival in both hosts (2,23,24). Consistent with this, SS cells were more resistant to culture in very low glucose media, a characteristic of the tsetse fly midgut environment (Fig. 1B). To explore the role that glucose depletion has on gene expression, the transcriptomes of LS parasites, SS parasites cultured in high glucose media, SS parasites cultured in very low glucose media, and PF parasites differentiated from very low glucose media were used to generate a principle response curve (PRC) (25) (S6 Fig). Overall, differentiation from LS to PF parasites yielded changes in expression of many life-stage specific genes (S1 Table). During the transition from LS to SS in the presence of glucose, there was a dramatic change in expression profiles. However, the removal of glucose in the SS life cycle stage only led to a minor alteration in gene expression (S6 Fig) with a total of 14 upregulated and 21 downregulated genes detected when SS parasites cultured in very low glucose were compared to those maintained in high glucose through differential gene expression analysis (S3 Table). Interestingly, three of the significantly regulated transcripts were also found to be among the top genes associated with progression from LS to PF (Table 1).

**Table 1.**
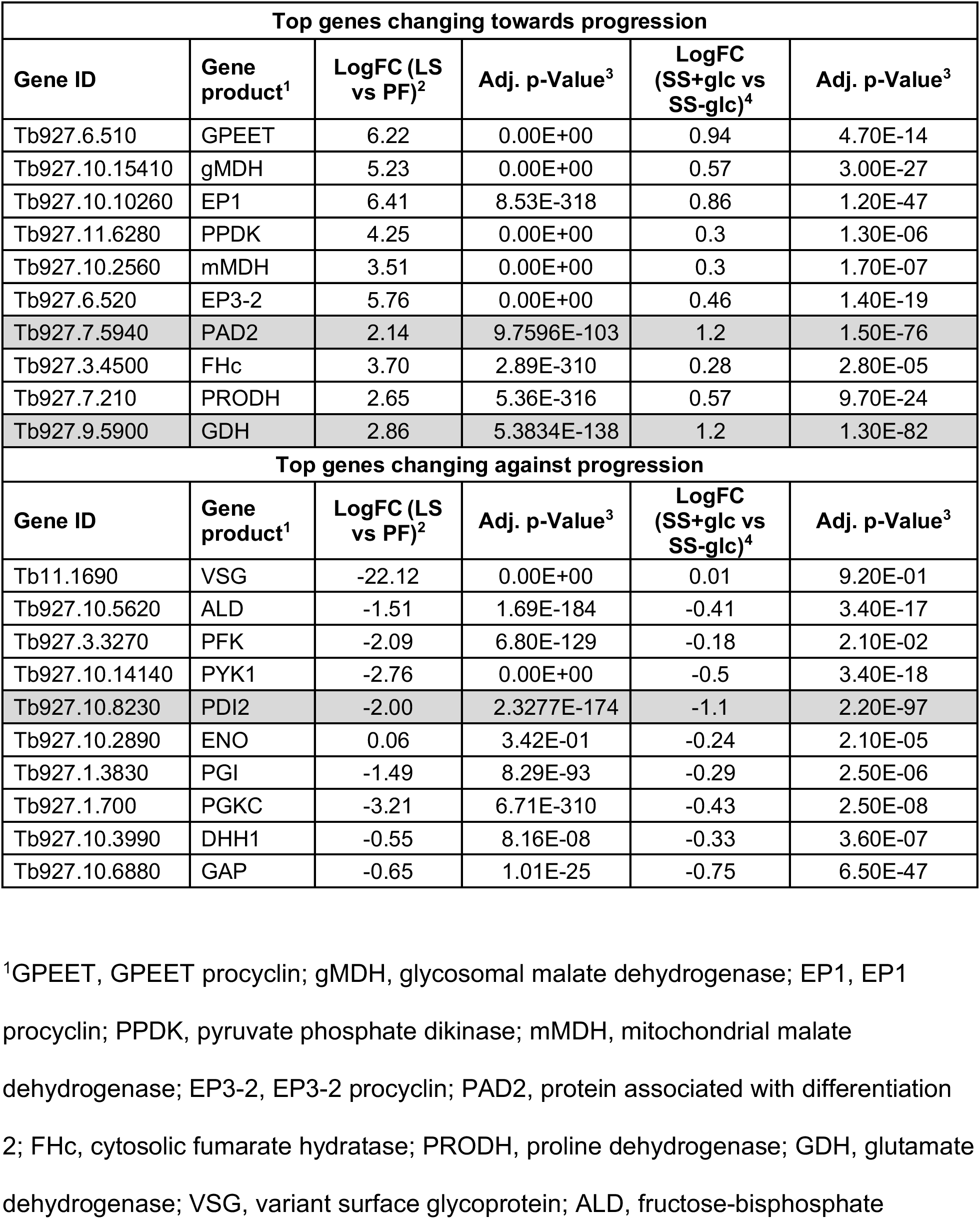
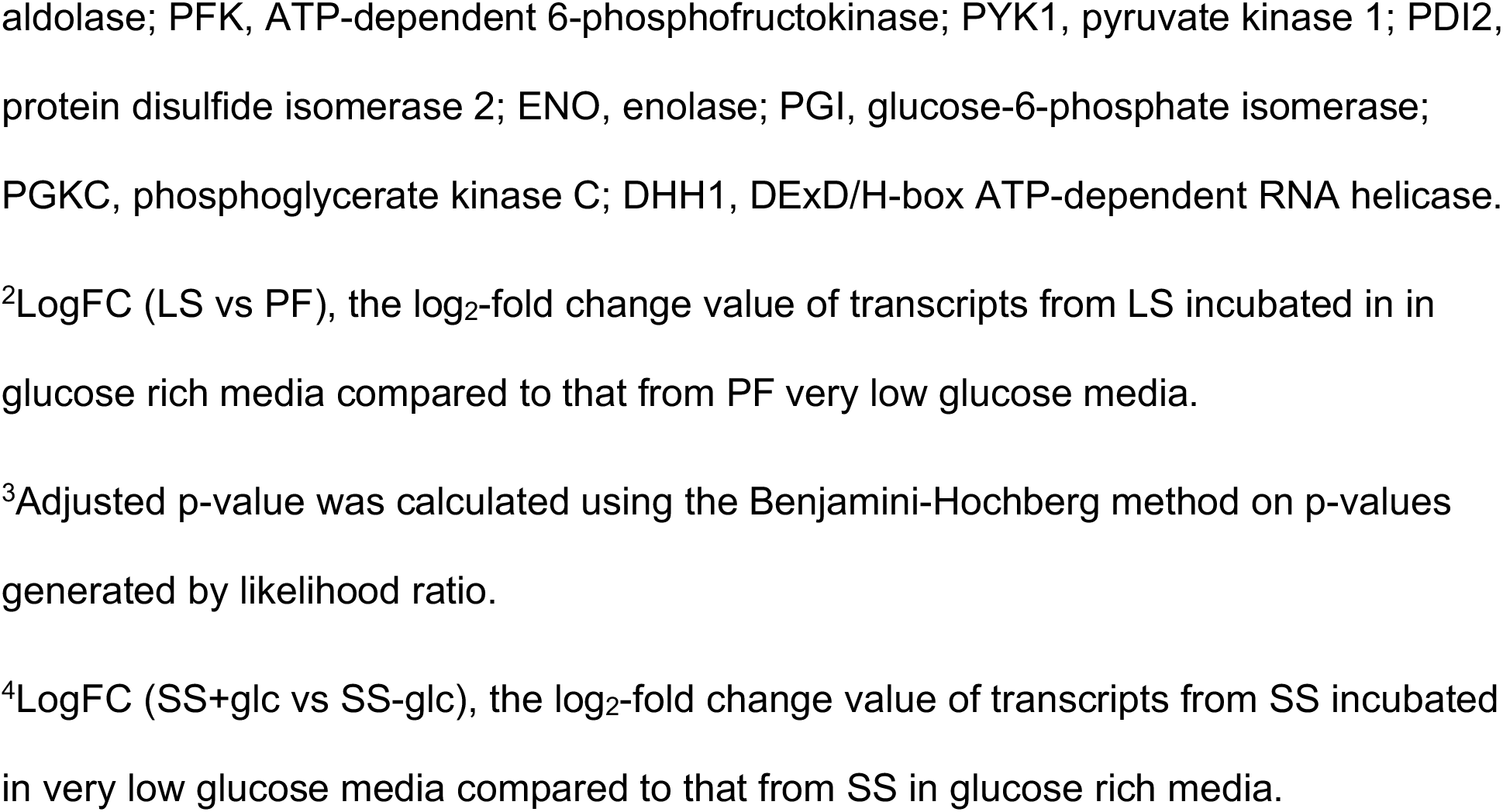
The fold-change and adjusted p-values from the SS (+glc +AA) vs. SS (-glc+AA) DGE comparison.

### Pleomorphic PF cell growth is inhibited by glucose

Differentiation of PF parasites from SS parasites by either citrate treatment (6 mM) or incubation in the near-absence of glucose yielded parasites with a growth deficiency upon culture in glucose-rich medium (Fig. 6A). These pleomorphic PF cells had average doubling times (between days 4 and 7) that were 4.3-fold and 2.9-fold greater, respectively, than the doubling time of parasites maintained in medium without glucose generated by the same differentiation means (Fig. 6A, compare filled symbols to open circles and asterisks). The growth defect in the presence of glucose persisted for several weeks, with cell division becoming nearly undetectable on day 12. However, cells continuously cultured in glucose-replete medium eventually (3-4 weeks) resumed growth (S4 Fig.).

**Fig 6.**
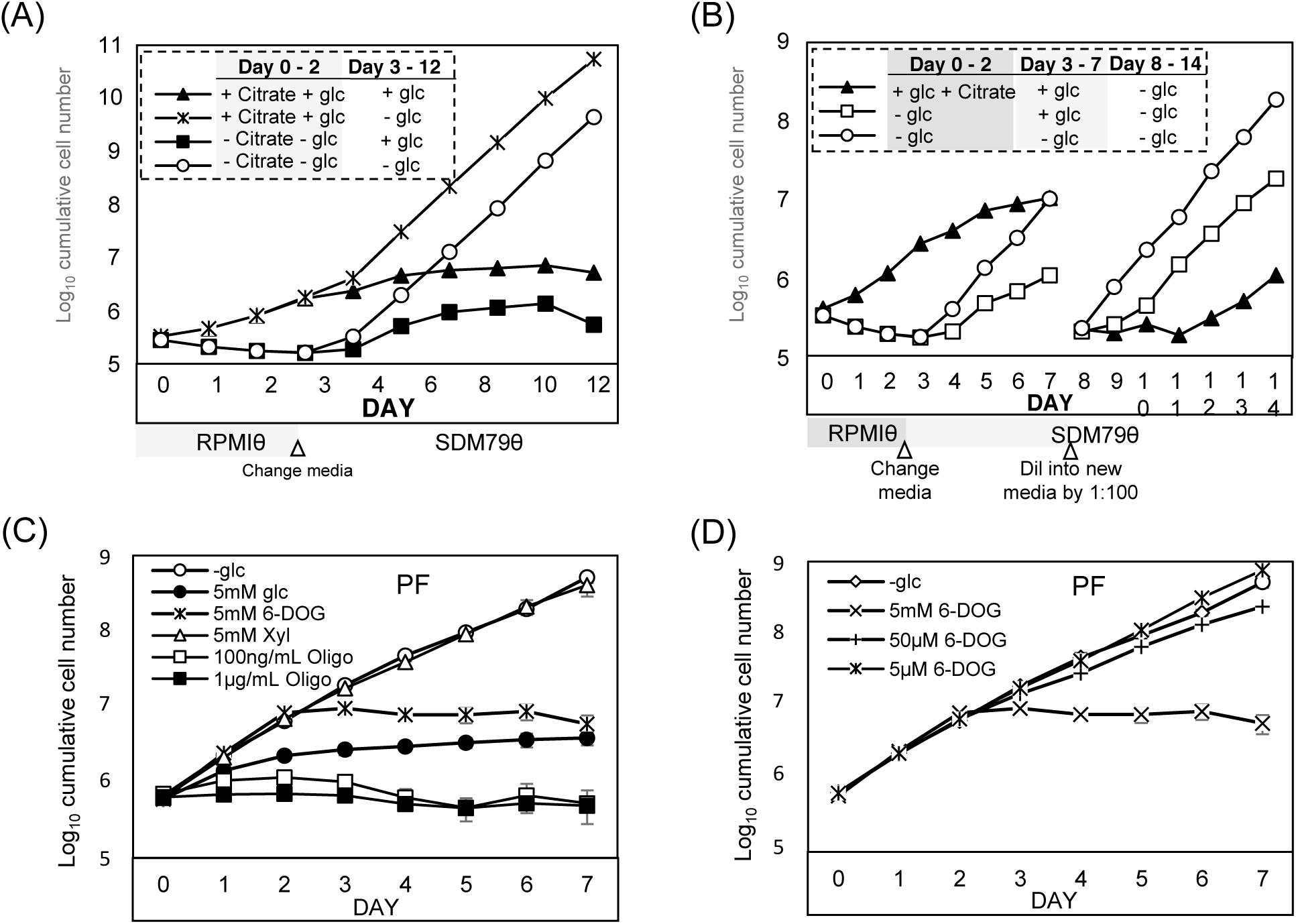
Pleomorphic PF cell growth is suppressed in the presence of glucose. (A) Comparison of growth of PF parasites differentiated from SS parasites by treatment with 6 mM citrate in the presence of glucose (5 mM) to those differentiated by glucose depletion. (B) Growth suppression was reversible by removal of glucose. SS parasites were cultured for two days in the presence of citrate (6 mM) or near-absence of glucose and then cultures were resuspended in glucose-replete or deficient SDM79θ for five days, prior to dilution in fresh medium on the eighth day of growth. (C) Growth of pleomorphic PF cells (differentiated for 7 days from SS) in SDM79θ with assorted sugars (6-DOG, 6-deoxy-glucose; xyl, xylose) or (D) in different concentrations of 6-DOG. Panel A and B are representative of at least duplicate assays, while in panel C the bars indicate standard deviation in triplicate assays.

The addition of 5 mM glucose to the environment slowed the growth of PF parasites previously differentiated by glucose deprivation when compared to similar cells maintained in very low glucose (Fig. 6B, open squares compared to open circles, day 3-7). This growth retardation was relieved within one day of glucose removal from the medium (open squares, day 8-14). While citrate-treated cells differentiated in the presence of glucose within one day (filled triangles), these PF parasites had a reduced growth rate when compared to cells differentiated by glucose depletion (open circles, day 3-7). This growth retardation was partially relieved after ∼3 days of culture in very low glucose (filled triangles, day 8-14) but the ∼30 hour doubling time still lagged behind the ∼18 hours doubling time of cells differentiated by glucose depletion (open squares, based on rates on days 11-14). Glucose was not alone in its ability to temper pleomorphic PF growth, as 5 mM 6-DOG, but not lower concentrations (Fig. 6D), was able to inhibit parasite growth even though it is not a metabolizable sugar (Fig. 6C, x symbol). Similarly, 2-DOG inhibited growth of pleomorphic PF parasites differentiated in the absence of glucose (not shown). Xylose had no impact on PF cell growth (Fig. 6C, open triangles).

### PF parasites differentiated in glucose depleted medium upregulate amino acids metabolism

Comparison of transcriptome profiles of LS parasites to PF parasites differentiated by culture of SS parasites in very low glucose (5 µM) by differential gene expression analyses revealed that a total of 1,310 transcripts were differentially-regulated in the PF cells, with 694 upregulated and 616 downregulated genes (false discovery rate (FDR) < 0.05, |logFC| ≥ 1.0, and counts per million (CPM) > 10; S1 Table for a complete list and verification of a subset by qRT-PCR). This result is in agreement with previous microarray studies which found that a similar number of total genes were differentially expressed between the LS and PF life stages (23,26).

To determine if the different cues used to trigger differentiation yielded distinct transcript profiles, the data sets for PF parasites differentiated by culture in very low glucose or by culture with CCA in the presence of glucose were compared to the transcriptome of LS parasites (23,26). Transcripts encoding proteins known to be enriched in BF parasites, like ESAG2, ESAG11, THT1-, and phosphoglycerate kinase (PGKC) were significantly downregulated in both PF transcriptomes, whereas those known to be enriched in PF parasites, including EP1, EP2, EP3, GPEET, trypanosome hexose transporter 2A (THT2A), purine nucleoside transporter NT10, and several proteins involved in electron transport chain and mitochondrial translation were significantly upregulated (Fig. 7A).

**Fig 7.**
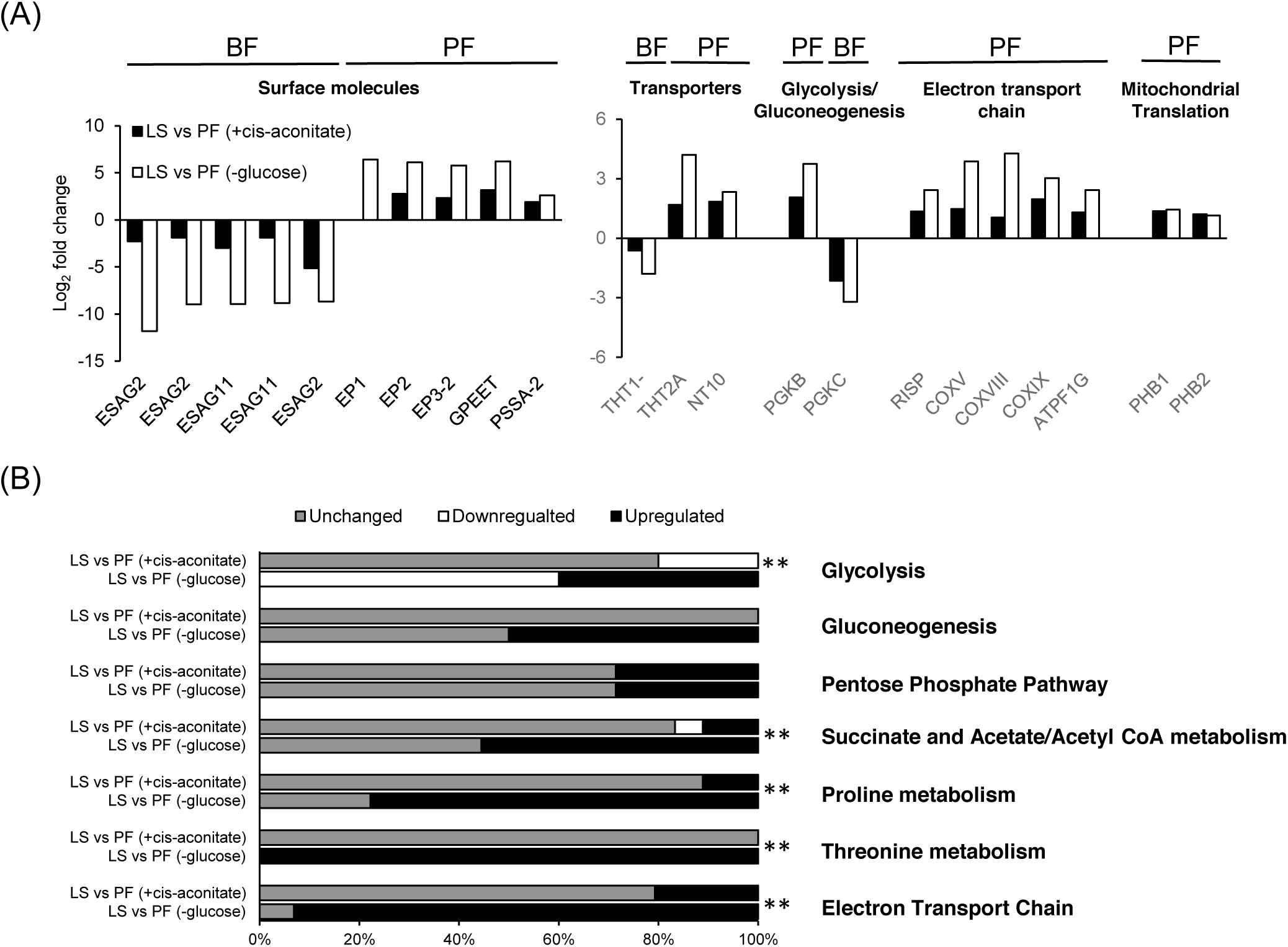
Evaluation of pleomorphic PF differentiated by different cues at the transcriptome level. (A) A comparison of the relative transcript abundance of PF parasites to LS parasites for a subsect of life stage-enriched genes. PF were generated by differentiation of SS parasites using either glucose depleted medium (-glc) or cis-aconitate treatment (+cis-aconitate). All transcripts included had >2-fold change and adjusted p-values < 0.05 except for THT1- in PF (+cis-aconitate), which was downregulated less than twofold with an adjusted p-value > 0.05. N/A, data not available. ESAG2, expression site-associated gene 2; ESAG11, expression site-associated gene 11; EP1, EP1 procyclin; EP2, EP2 procyclin; EP3-2, EP3-2 procyclin; GPEET, GPEET procyclin; PSSA-2, procyclic form surface phosphoprotein; THT1-, glucose transporter 1B; THT2A, glucose transporter 2A; NT10, purine nucleoside transporter 10; PGKB, phosphoglycerate kinase B; PGKC, phosphoglycerate kinase C; RISP, rieske iron-sulfur protein; COXV, cytochrome oxidase subunit V; COXVIII, cytochrome oxidase subunit VIII; COXIX, cytochrome oxidase subunit IX; ATPF1G, ATP synthase F1 subunit gamma protein; PHB1, prohibitin 1, PHB2, prohibitin 2. (B) The percentage of transcripts in major metabolic pathways (as identified by GO term definition and confirmed in TrypanoCyc (57)) that are regulated in PF differentiated by cis-aconitate or glucose depletion compared to LS. For glycolysis and gluconeogenesis, only enzymes that participate in one of the reactions were included. Upregulated transcripts are those that increased > 2-fold and had adjusted p-values < 0.05 in PF compared to LS, while downregulated transcripts were decreased > 2-fold with adjusted p-values < 0.05. Chi-squared goodness of fit tests were performed for genes in each pathway to determine whether the proportions of regulated genes between PF differentiated by cis-aconitate and glucose depletion were different. ** indicates p-value < 0.01, suggesting transcripts involved in the pathway are differentially regulated in PFs differentiated by the two cues.

While these life-stage-specific marker genes were similarly regulated, differences in the regulation of genes involved in major metabolic pathways were observed. Genes required for glycolysis, the primary catabolic pathway used for ATP production in BF parasites, were largely downregulated (60% of the known or predicted genes in the pathway) in PF parasites differentiated by culture in glucose depleted medium, while PF parasites differentiated by the presence of cis-aconitate had 20% of the known or predicted genes in the pathway downregulated (Fig. 7B, S2 Table). The percentage of upregulated genes involved in gluconeogenesis was higher in PF parasites differentiated by glucose depletion (50%) than those differentiated by cis-aconitate, where the expression of genes in the pathway remained largely unchanged (Fig. 7B, S2 Table). Similarly, the percentage of upregulated genes known to be required for succinate and acetate/acetyl-CoA metabolism were higher in PF parasites generated by glucose depletion (56% of known genes involved) than in cis-aconitate differentiated PF parasites (11%).

A larger difference was noted for changes in expression of genes in pathways associated with amino acid metabolism. While the cis-aconitate differentiated PF parasites showed upregulation in several transcripts involved in proline catabolism and electron transport chain (such as succinyl-CoA synthetase alpha subunit (Tb927.3.2230), succinate dehydrogenase assembly factor 2 (SDHAF2, Tb927.6.2510), and the mitochondrial precursor of Rieske iron-sulfur protein (RISP, Tb927.9.14160), to name a few, the majority of transcripts in the pathways were unchanged (Fig. 7B, S2 Table). By comparison, expression of most of the known genes involved in proline, threonine, and electron transporter chain (∼78%, ∼100%, and ∼93%, respectively) were upregulated in PF parasites generated by GRS differentiation.

## Discussion

Cellular mechanisms for adaptation to environmental nutrient availability are found in organisms as divergent as microbes and humans. These mechanisms typically initiate upregulation of alternative metabolic pathways, reduction of macromolecule synthesis in response to limited nutrients, and enhancement of the expression of genes that may improve organism dispersal to ultimately lead to colonization of new, nutrient-rich environments. Additional responses include those that trigger development into quiescent or environmental-resistant forms that can tolerate exposure to hostile environments.

The rapid depletion of glucose from a tsetse fly bloodmeal, as described by Vickerman (11), suggests that glucose abundance could serve as a cue for developmental programing. However, the rate of GRS differentiation is, by itself, likely too slow to be biologically relevant. It is interesting, however, that combination with other treatments known to influence differentiation (but that are by themselves insufficient to trigger differentiation) potentiated the rate of differentiation to levels similar to those observed with described cues. A temporary cold shock treatment (9) combined with culture in very low glucose led to detectable EP procyclin protein expression after three hours and outgrowth within one day. This was a notable reduction from the three-day period required when parasites were grown in minimal glucose alone (Fig. 3). Cold shock treatment also upregulates the carboxylate transporter PAD2 and is therefore important for sensitizing SS parasites to physiologically-relevant levels of CCA (3,4). Interestingly, glucose depletion also upregulated PAD2 expression. Given that glucose is precipitously depleted from the blood meal (11), the upregulation of PAD2 in response to glucose depletion may serve as an additional or alternative means of increasing SS parasite sensitivity to CCA.

The differentiation of SS parasites in response to reduced glucose availability required inclusion of amino acids in the medium and yielded PF parasites with transcriptome-wide alteration of the expression of major metabolic pathway genes, a pattern not found in PF parasites differentiated through cis-aconitate treatment. This difference suggests that glucose availability may influence activation of metabolic pathways required for different developmental stages. To assess if the gene expression pattern was uniquely associated with glucose depletion, the transcriptome changes in response to glucose depletion were compared to transcriptional data generated from analysis of nine other transcriptomes (S2 Table) (17,26-33). The analyses include transcriptomes from pleomorphic Antat1.1 LS and PF and pleomorphic TREU927 LS and PF generated from mouse-derived SS parasites with 6 mM of cis-aconitate, monomorphic Lister 427 BF and lab-adapted PF parasites, and monomorphic BF (Lister 427) parasites with RBP10 knocked down. Five of these with accessible statistical analysis were chosen as representatives and differences and similarities to the transcriptome generated in this study were compared (S5 Fig). The cells used in these treatments were maintained in media that contained at least 5 mM glucose (17,23,27,29,32).

While PF cells generated in these studies generally downregulated glycolytic genes and upregulated amino acid metabolism genes, the proportion of the genes differentially regulated in these categories was significantly different from the regulation generated by glucose depletion (S5 Fig). Interestingly, the upregulation of the two hexokinase genes was only observed when PF were maintained in media with very low glucose levels or when the cells were supplemented with glycerol (S2 Table, Qiu, Queiroz, and Naguleswaran columns). While the trend was not statistically significant, the overall regulation of the glycolytic and amino acid metabolic pathways in these three PF lines was highly similar. This similarity may not be a coincidence but may reflect the influence of glucose and glycerol on a regulatory pathway. Consistent with this, inclusion of glucose or glycerol in growth medium have been shown to have the opposite impact on *GPEET* transcript levels, a marker of early PF parasites (19).

In *T. brucei*, the transcriptional regulation largely occurs at the post-transcriptional level, a process that frequently involves the action of RNA binding proteins (34). One such RNA binding protein, RBP10, has been shown to play a major role during the transition of parasites from blood to insect stages (17,35). As an RNA binding protein that is exclusively expressed in BF parasites, RBP10 influences gene expression at both transcript and protein levels. Knocking down RBP10 in BF parasites has been found to trigger a series of developmental changes that eventually give rise to PF parasites (35). Consistent with a role in regulating development, RBP10 transcript levels were found to be downregulated in most of the PF transcriptomes compared here, including the one derived from glucose depletion in this study (S2 Table). However, the massive alteration of transcript abundance for genes involved in energy metabolism was also not observed in RBP10 knockdown BF cells (S5 Fig; S2 Table). This incomplete overlap of transcript profiles suggests that glucose signaling may influence transcript level through pathways yet to be discovered.

GRS differentiation includes a nutrient adaptation mechanism that responds to glucose, either as a result of the perception of a signal through glucose metabolism or by signaling through a glucose-specific receptor-mediated pathway. The observation that GRS differentiation in SS parasites was inhibited by low concentrations of 6-DOG, a glucose analog that is not a substrate for glycolysis, suggests that *T. brucei* perceives the presence of the hexose through a glycolysis-independent pathway. In other eukaryotes, glucose receptor-mediated signaling is known to influence cellular responses to extracellular glucose (36,37). The identity of potential glucose receptors that would participate in this sort of response in *T. brucei* is unknown, but it is possible that one or more of the trypanosome hexose transporter family members, or one of the hexokinases, could serve this role, given that they have similar *K*_M_ values (52 μM for the trypanosome hexose transporter 2 and 90 μM for the trypanosome hexokinase 1) to the concentration of glucose required to inhibit GRS differentiation (50 μM) (38,39). However, the involvement of glucose metabolism in cellular development and nutrient adaptation cannot be completely ruled out. Treatment of monomorphic BF with phloretin, a glucose uptake inhibitor, has been shown to trigger a differentiation-like transcriptome change in the parasites (16).

The pleomorphic PF parasites generated by the GRS differentiation offer additional evidence for two pathways to sense glucose. First, 6-DOG inhibited growth of these PF cells, suggesting metabolism was connected to the observed growth repression. However, the onset of growth inhibition by addition of glucose to the medium was more rapid than that observed using 6-DOG (Fig. 6C), possibly because it activated both the glycolysis-dependent and the glucose receptor-mediated pathways simultaneously, while 6-DOG could only initiate the receptor-mediated pathway. Alternatively, glucose uptake may have been more efficient which could impact the activity of an intracellular glucose binding protein-based response.

While LS parasites are dependent on host glucose for ATP production, PF parasites have a more dynamic metabolism and are capable of utilizing either amino acids or sugars as carbon sources. Laboratory-adapted PF parasites grown in glucose-rich culture media preferentially metabolize glucose with coincident downregulation of proline consumption (40,41). PF parasites generated by culture of SS forms in reduced glucose media responded to the sugar differently, by displaying growth inhibition (Fig. 6). This was similar to glucose-induced growth inhibition noted in PF parasites isolated from tsetse fly midguts (42), which presumably occurred because these parasites had been adapted to the amino acid-rich fly gut environment (43). Additionally, the negative impact of glucose on parasite growth may explain why trypanosome infections are more readily established in blood-starved tsetse flies than in fed insects (44,45).

In conclusion, the glucose signaling pathway in the African trypanosome may not be directly connected to glycolysis, but the pathway does play an important role in the metabolic adaptation of the parasite (Fig. 7, S5 Fig, S2 Table), which may in turn influence other aspects of developmental differentiation through known pathways.

## Methods

### Ethics Statement

All procedures were carried out in accordance with the PHS Policy on the Care and Use of Laboratory Animals and in accord with the CU PHS Assurance Number A3737-01 under the approval of the Clemson University Institutional Animal Care and Use Committee (IACUC). Clemson University animal research and teaching facilities and programs are registered by USDA, Animal Care (AC) [Registration Number 56-R-0002] and animal research programs and facilities have full accreditation from the Association for Assessment and Accreditation of Laboratory Animal Care, International (AAALAC). Euthanasia by heavy anesthetization followed by bilateral pneumothorax was used based on recommendations from the AVAM Guidelines for the Euthanasia of Animals.

### Trypanosomes and cell culture conditions

*Trypanosoma brucei brucei* 427 BF parasites, a representative monomorphic strain, were cultured as described in HMI-9 medium (46). LS and SS *T. brucei* AnTat1.1 were isolated from infected Swiss Webster mice 3-4 or 6-7 days after infection, respectively, by preparation of buffy coats and purification through DEAE chromatography. Cells were maintained in HMI-9 medium (with LS density kept under 5 × 10^5^ cells/mL) for one day before use.

To assess trypanosome responses to environmental manipulations, a BF medium with very low glucose, RPMIθ, was made by modifying HMI-9 (46) with additional features adapted from (47) including replacement of IMDM with glucose-free RPMI that has been buffered with HEPES to pH 7.4, elimination of SerumPlus and use of dialyzed FBS (10% f.c.). A PF medium with very low glucose, SDM79θ, was generated by eliminating glucose from SDM79 (48) using dialyzed FBS (10% f.c.) in lieu of standard serum. Both medium have a final glucose concentration of ∼5 μM. To score the impact of culturing under extremely low glucose conditions, parasites were washed three times in warmed PBS to eliminate residual glucose before resuspension at ∼5 × 10^5^ cells/mL in prewarmed (at 27°C) SDM79θ with or without additional carbon sources or glucose analogs added. Cultures were incubated at 27°C or 37°C in 5% CO_2_ with their medium changed every two days. Cell numbers were scored daily during the first week and every-other-day in the second week and cell viability determined by propidium iodide (PI) staining (0.5 µg/ml final concentration) followed by flow cytometry on an Accuri BD flow cytometer (BD Biosciences, San Jose, CA).

### Glucose measurements

To assay the glucose concentration in the parasite growth medium, an Amplex™ Red Glucose/Glucose Oxidase Assay Kit (Invitrogen, Carlsbad CA) was used. All time points were tested in triplicate, with parasites removed by centrifugation (16,000 × g, 2 min) and supernatant tested according to the protocol provided by the manufacturer.

### RNA analysis

To assess the consequence of glucose availability on steady-state transcript abundance, RNAseq was performed on LS cells isolated from rodents and cultured in HMI9 for one day, SS cells incubated for one day in RPMIθ with or without added glucose (5 mM) supplemented with or without proline and threonine, and PF cells differentiated from SS in very low glucose. Three biological replicates of each treatment were conducted and RNA was isolated using an Aurum Total RNA Mini Kit (Bio-Rad). Libraries were constructed using the TruSeq Stranded mRNA Library Prep Kit (Illumina, San Diego, CA USA). Quality metrics were analyzed for all samples using FastQC (http://www.bioinformatics.babraham.ac.uk/projects/fastqc). Trimming of low quality bases and adapter sequences was performed using trimmomatic (49). Trimmed reads were aligned using gsnap to the TriTrypDB-28_TbruceiTREU927 genome, minus the 11 bin scaffold/chromosome. Subread’s featureCounts was used to identify reads uniquely assigned to known genes in the correct direction of transcription (50). Raw read counts were then used as input to edgeR for differential gene expression (DGE) analysis using generalized linear models (GLM) (51,52). Genes with low coverage across all samples were filtered out, and library sizes were normalized using the trimmed mean of M-values method. Sample comparisons were set up as likelihood ratio tests, and genes having a FDR of < 0.05, |logFC| ≥ 1.0, and CPM >10 were considered to have significant expression abundance changes.

To verify the quality of RNAseq, qRT-PCR was performed using a Verso 1-step RT-qPCR Kit (ThermoFisher) in a CFX96 Touch™ Real-Time PCR Detection System (Bio-Rad) for selected transcripts. Ct values of transcripts were used to solve relative expression by the comparative Ct (2^-ΔΔCT^) method using the expression of the telomerase reverse transcriptase (TERT) gene or 60S ribosomal protein L10a (RPL10A) as reference as described (16,53,54). Primers for transcripts were designed through Genscript Real-time PCR (TaqMan) Primer Design (https://www.genscript.com/tools/real-time-pcr-tagman-primer-design-tool) and confirmed by blasting against the *Trypanosoma brucei brucei* TREU927 transcriptome database on TritrypDB (http://tritrypdb.org/tritrypdb/). A list of primers used in qRT-PCR can be found in S4 table.

### Flow cytometry and epifluorescence microscopy

For analysis of surface molecule expression of parasites by flow cytometry, antibody staining was performed using protocols modified from (55). Briefly, 0.5-2 × 10^6^ cells were washed in PBS and fixed in 4% formaldehyde/0.05% glutaraldehyde for 1 hour at 4°C. Cells then incubated in blocking solution (2% BSA in PBS) for 1 hour before application of FITC-conjugated EP procyclin antibody (monoclonal antibody TBRP1/247, Cedarlane Laboratories, 1:1000) or PAD1 antibody (a generous gift of Dr. Keith Matthews, University of Edinburgh, 1:1000). Cells were washed three times in PBS before addition of Alexa Fluor 488-conjugated goat anti-mouse (1:1000) or goat anti-rabbit secondary antibody (ThermoFisher Scientific, 1:1000) prior to analysis by flow cytometry.

Immunofluorescence assays were performed using a protocol modified from (56). Parasites (1 × 10^6^ cells) were harvested (800 × *g*, 8 min), washed with PBS, and then fixed in 4% formaldehyde in PBS for 30 minutes at 4°C. Cells were washed with PBS, allowed to settle on poly-lysine coated slides, and permeabilized with 0.1% Triton X-100 in PBS for 30 min. After being washed in PBS, cells were incubated in blocking solution (10% normal goat serum and 0.1 % Triton X-100 in PBS) for 1 hour at room temperature, followed by addition of the FITC conjugated EP procyclin antibody diluted at 1:100 or PAD1 antibody diluted at 1:100 in blocking solution. Primary antibodies were detected with Alexa Fluor 488-conjugated antiserum diluted at 1:1000. Vectashield mounting medium with DAPI was applied for the detection of nucleus and kinetoplast DNA. Cells were visualized on a Zeiss Axiovert 200M using Axiovision software version 4.6.3 for image analysis.

## Acknowledgements

The authors would like to thank Sarah Grace McAlpine for her technical assistance and Drs. Kimberly Paul and Meredith Morris for their comments on the manuscript. We also thank Dr. Keith Matthews for generously supplying the PAD1 antisera.

## Abbreviations

BF: bloodstream form *T. brucei*
GDP: glucose-dependent pathway
GIP: glucose-independent pathway
IF: immunofluorescence
LS: long slender bloodstream form *T. brucei*
PF: procyclic form *T. brucei*
RPMIθ: a BF medium with minimal glucose
SDM79θ: a PF medium with minimal glucose
SS: short stumpy bloodstream form *T. brucei*

## Supporting information

**S1 Fig. Monomorphic BF do not differentiate to SS in RPMIθ.** In triplicate, monomorphic BF 427 parasites were washed, resuspended in RPMIθ, supplemented with or without glucose or glycerol (5 mM) and cultured for two days. Parasites were then collected, washed, and resuspended in RPMIθ supplemented with glucose (filled circles) or in PF medium lacking glucose (SDM79θ, ∼5 µM glucose) supplemented with or without additional glucose (open symbols).

**S2 Fig. Alternative sugar analogs do not support growth of glycolysis-dependent LS parasites**. LS parasites (3 × 10^4^/mL) were extensively washed and resuspended in very low glucose medium, RPMIθ, supplemented with the indicated compounds (glc, glucose; xyl, xylose; 2-DOG, 2-deoxyglucose; 6-DOG, 6-deoxyglucose) at 5 mM and growth monitored as described.

**S3 Fig. Outgrowth of SS parasites seeded in different concentrations of glucose.** SS parasites were washed extensively and resuspended in RPMIθ supplemented with varied glucose (5 µM- 5 mM) for two days prior to transfer to SDM79θ and growth monitored.

**S4 Fig. Long-term growth of pleomorphic PF parasites differentiated by glucose depletion (-glc) or addition of 6 mM citrate (cit) in the presence or absence of glucose (5 mM).** Growth assays were conducted using cells that were harvested seven days after differentiation. For cells that stopped growing in glucose-rich medium (+glc), medium was replaced every 3-4 days to maintain the glucose concentration.

**S5 Fig. Evaluation of transcriptomes from pleomorphic PF parasites differentiated with different cues.** The percentage of transcripts in major metabolic pathways (as identified by GO term definition and confirmed in TrypanoCyc (57)) that were regulated in PF parasites differentiated by cis-aconitate or glucose depletion compared to LS parasites. For glycolysis and gluconeogenesis, only enzymes that participate in one of the reactions were included. Chi-squared goodness of fit tests were performed for genes in each pathway to compare the regulation patterns of GRS differentiated PF parasites to those from other treatments. ** indicates p-value < 0.01, suggesting transcripts involved in the pathway are differentially regulated in PF parasites differentiated by the two cues.

**S6 Fig. SS parasites are largely preadapted at the transcript level to become PF parasites.** A principle responsive curve (PRC) that illustrates the global transcriptome progression from LS to PF parasites (solid dark grey line) through transitions that include SS parasites in glucose-replete and very low glucose media. The explanatory variable (time) was based on when each transition occurred and were as follows: Day 0, LS form with glucose (LS+glc); Day 3, SS form maintained with glucose (SS+glc); Day 4, SS form (from Day 3) cultured in the absence of glucose for one day; Day 10, PF without glucose (PF-glc). PRC was designed such that the comparison to baseline (Day 0, LS+glc, dashed grey line) was treated as repeated measurements for each time point. The unscaled variable coefficients (Species Scores) have been depicted as a sorted line stack graph on the right. The top transcripts with annotation that changed towards (dark grey) and against (light grey) the LS to PF parasite progression have been identified on the line stack graph.

**S1 Table. A complete list of DEG in PF parasites differentiated from glucose depletion compared to LS parasites and validation by qRT-PCR.**

**S2 Table. A list of genes used in comparative gene expression analysis of essential metabolic pathways.**

**S3 Table. A complete list of DEG in SS incubated in very low glucose medium compared to SS in glucose rich medium and validation by qRT-PCR.**

**S4 Table. A complete list of qRT-PCR primers used in this study.**

## References

1. Reuner, B., Vassella, E., Yutzy, B., and Boshart, M. (1997) Cell density triggers slender to stumpy differentiation of *Trypanosoma brucei* bloodstream forms in culture. Mol Biochem Parasitol 90, 269–280

2. Rico, E., Rojas, F., Mony, B. M., Szoor, B., Macgregor, P., and Matthews, K. R. (2013) Bloodstream form pre-adaptation to the tsetse fly in *Trypanosoma brucei*. Front Cell Infect Microbiol 3, 78

3. Engstler, M., and Boshart, M. (2004) Cold shock and regulation of surface protein trafficking convey sensitization to inducers of stage differentiation in *Trypanosoma brucei*. Genes Dev 18, 2798–2811

4. Dean, S., Marchetti, R., Kirk, K., and Matthews, K. R. (2009) A surface transporter family conveys the trypanosome differentiation signal. Nature 459, 213–217

5. Hunt, M., Brun, R., and Kohler, P. (1994) Studies on compounds promoting the in vitro transformation of *Trypanosoma brucei* from bloodstream to procyclic forms. Parasitol Res 80, 600–606

6. Rolin, S., Hancocq-Quertier, J., Paturiaux-Hanocq, F., Nolan, D. P., and Pays, E. (1998) Mild acid stress as a differentiation trigger in *Trypanosoma brucei*. Mol Biochem Parasitol 93, 251–262

7. Yabu, Y., and Takayanagi, T. (1988) Trypsin-stimulated transformation of *Trypanosoma brucei* gambiense bloodstream forms to procyclic forms in vitro. Parasitol Res 74, 501–506

8. Sbicego, S., Vassella, E., Kurath, U., Blum, B., and Roditi, I. (1999) The use of transgenic *Trypanosoma brucei* to identify compounds inducing the differentiation of bloodstream forms to procyclic forms. Mol Biochem Parasitol 104, 311–322

9. Szoor, B., Dyer, N. A., Ruberto, I., Acosta-Serrano, A., and Matthews, K. R. (2013) Independent pathways can transduce the life-cycle differentiation signal in *Trypanosoma brucei*. PLoS Pathog 9, e1003689

10. Szoor, B., Ruberto, I., Burchmore, R., and Matthews, K. R. (2010) A novel phosphatase cascade regulates differentiation in *Trypanosoma brucei* via a glycosomal signaling pathway. Genes Dev 24, 1306–1316

11. Vickerman, K. (1985) Developmental cycles and biology of pathogenic trypanosomes. Br. Med. Bull. 41, 105–114.

12. Mantilla, B. S., Marchese, L., Casas-Sanchez, A., Dyer, N. A., Ejeh, N., Biran, M., Bringaud, F., Lehane, M. J., Acosta-Serrano, A., and Silber, A. M. (2017) Proline metabolism is essential for *Trypanosoma brucei brucei* survival in the tsetse vector. PLoS Pathog 13, e1006158

13. Millerioux, Y., Ebikeme, C., Biran, M., Morand, P., Bouyssou, G., Vincent, I. M., Mazet, M., Riviere, L., Franconi, J. M., Burchmore, R. J., Moreau, P., Barrett, M. P., and Bringaud, F. (2013) The threonine degradation pathway of the *Trypanosoma brucei* procyclic form: the main carbon source for lipid biosynthesis is under metabolic control. Mol Microbiol 90, 114–129

14. Smith, T. K., Bringaud, F., Nolan, D. P., and Figueiredo, L. M. (2017) Metabolic reprogramming during the *Trypanosoma brucei* life cycle. F1000Res 6

15. Milne, K. G., Prescott, A. R., and Ferguson, M. A. (1998) Transformation of monomorphic *Trypanosoma brucei* bloodstream form trypomastigotes into procyclic forms at 37 degrees C by removing glucose from the culture medium. Mol Biochem Parasitol 94, 99–112.

16. Haanstra, J. R., Kerkhoven, E. J., van Tuijl, A., Blits, M., Wurst, M., van Nuland, R., Albert, M. A., Michels, P. A., Bouwman, J., Clayton, C., Westerhoff, H. V., and Bakker, B. M. (2011) A domino effect in drug action: from metabolic assault towards parasite differentiation. Mol Microbiol 79, 94–108

17. Wurst, M., Seliger, B., Jha, B. A., Klein, C., Queiroz, R., and Clayton, C. (2012) Expression of the RNA recognition motif protein RBP10 promotes a bloodstream-form transcript pattern in Trypanosoma brucei. Mol Microbiol 83, 1048–1063

18. Morris, J. C., Wang, Z., Drew, M. E., and Englund, P. T. (2002) Glycolysis modulates trypanosome glycoprotein expression as revealed by an RNAi library. EMBO J. 21, 4429–4438

19. Vassella, E., Probst, M., Schneider, A., Studer, E., Renggli, C. K., and Roditi, I. (2004) Expression of a major surface protein of *Trypanosoma brucei* insect forms is controlled by the activity of mitochondrial enzymes. Mol Biol Cell 15, 3986–3993

20. Feagin, J. E., Jasmer, D. P., and Stuart, K. (1986) Differential mitochondrial gene expression between slender and stumpy bloodforms of *Trypanosoma brucei*. Mol Biochem Parasitol 20, 207–214

21. Bienen, E. J., Saric, M., Pollakis, G., Grady, R. W., and Clarkson, A. B., Jr. (1991) Mitochondrial development in *Trypanosoma brucei brucei* transitional bloodstream forms. Mol Biochem Parasitol 45, 185–192

22. Gancedo, J. M. (2008) The early steps of glucose signalling in yeast. FEMS Microbiol Rev 32, 673–704

23. Kabani, S., Fenn, K., Ross, A., Ivens, A., Smith, T. K., Ghazal, P., and Matthews, K. (2009) Genome-wide expression profiling of in vivo-derived bloodstream parasite stages and dynamic analysis of mRNA alterations during synchronous differentiation in Trypanosoma brucei. BMC Genomics 10, 427

24. MacGregor, P., Szoor, B., Savill, N. J., and Matthews, K. R. (2012) Trypanosomal immune evasion, chronicity and transmission: an elegant balancing act. Nat Rev Microbiol 10, 431–438

25. Van den Brink, P. J., and Braak, C. J. F. T. (1999) Analysis of time-dependent multivariate responses of biological community to stress. Environ Tox Chem 18, 10

26. Queiroz, R., Benz, C., Fellenberg, K., Hoheisel, J. D., and Clayton, C. (2009) Transcriptome analysis of differentiating trypanosomes reveals the existence of multiple post-transcriptional regulons. BMC Genomics 10, 495

27. Jensen, B. C., Sivam, D., Kifer, C. T., Myler, P. J., and Parsons, M. (2009) Widespread variation in transcript abundance within and across developmental stages of Trypanosoma brucei. BMC Genomics 10, 482

28. Nilsson, D., Gunasekera, K., Mani, J., Osteras, M., Farinelli, L., Baerlocher, L., Roditi, I., and Ochsenreiter, T. (2010) Spliced leader trapping reveals widespread alternative splicing patterns in the highly dynamic transcriptome of *Trypanosoma brucei*. PLoS Pathog 6, e1001037

29. Siegel, T. N., Hekstra, D. R., Wang, X., Dewell, S., and Cross, G. A. (2010) Genome-wide analysis of mRNA abundance in two life-cycle stages of *Trypanosoma brucei* and identification of splicing and polyadenylation sites. Nucleic Acids Res 38, 4946–4957

30. Jensen, B. C., Ramasamy, G., Vasconcelos, E. J., Ingolia, N. T., Myler, P. J., and Parsons, M. (2014) Extensive stage-regulation of translation revealed by ribosome profiling of *Trypanosoma brucei*. BMC Genomics 15, 911

31. Vasquez, J. J., Hon, C. C., Vanselow, J. T., Schlosser, A., and Siegel, T. N. (2014) Comparative ribosome profiling reveals extensive translational complexity in different *Trypanosoma brucei* life cycle stages. Nucleic Acids Res 42, 3623–3637

32. Mugo, E., Egler, F., and Clayton, C. (2017) Conversion of procyclic-form Trypanosoma brucei to the bloodstream form by transient expression of RBP10. Mol Biochem Parasitol 216, 49–51

33. Naguleswaran, A., Doiron, N., and Roditi, I. (2018) RNA-Seq analysis validates the use of culture-derived *Trypanosoma brucei* and provides new markers for mammalian and insect life-cycle stages. BMC Genomics 19, 227

34. Kolev, N. G., Ullu, E., and Tschudi, C. (2014) The emerging role of RNA-binding proteins in the life cycle of *Trypanosoma brucei*. Cell Microbiol 16, 482–489

35. Mugo, E., and Clayton, C. (2017) Expression of the RNA-binding protein RBP10 promotes the bloodstream-form differentiation state in *Trypanosoma brucei*. PLoS Pathog 13, e1006560

36. Lemaire, K., Van de Velde, S., Van Dijck, P., and Thevelein, J. M. (2004) Glucose and sucrose act as agonist and mannose as antagonist ligands of the G protein-coupled receptor Gpr1 in the yeast *Saccharomyces cerevisiae*. Mol Cell 16, 293–299

37. Moriya, H., and Johnston, M. (2004) Glucose sensing and signaling in *Saccharomyces cerevisiae* through the Rgt2 glucose sensor and casein kinase I. Proc Natl Acad Sci U S A 101, 1572–1577

38. Barrett, M. P., Tetaud, E., Seyfang, A., Bringaud, F., and Baltz, T. (1998) Trypanosome glucose transporters. Mol Biochem Parasitol 91, 195–205.

39. Morris, M. T., DeBruin, C., Yang, Z., Chambers, J. W., Smith, K. S., and Morris, J. C. (2006) Activity of a second *Trypanosoma brucei* hexokinase is controlled by an 18-amino-acid C-terminal tail. Eukaryot Cell 5, 2014–2023

40. Lamour, N., Riviere, L., Coustou, V., Coombs, G. H., Barrett, M. P., and Bringaud, F. (2005) Proline metabolism in procyclic *Trypanosoma brucei* is down-regulated in the presence of glucose. J Biol Chem 280, 11902–11910

41. Coustou, V., Biran, M., Breton, M., Guegan, F., Riviere, L., Plazolles, N., Nolan, D., Barrett, M. P., Franconi, J. M., and Bringaud, F. (2008) Glucose-induced remodeling of intermediary and energy metabolism in procyclic *Trypanosoma brucei*. J Biol Chem 283, 16342–16354

42. van Grinsven, K. W., Van Den Abbeele, J., Van den Bossche, P., van Hellemond, J. J., and Tielens, A. G. (2009) Adaptations in the glucose metabolism of procyclic *Trypanosoma brucei* isolates from tsetse flies and during differentiation of bloodstream forms. Eukaryot Cell 8, 1307–1311

43. Bursell, E., Billing, K. J., Hargrove, J. W., McCabe, C. T., and Slack, E. (1973) The supply of substrates to the flight muscle of tsetse flies. Trans R Soc Trop Med Hyg 67, 296

44. Aksoy, S., Gibson, W. C., and Lehane, M. J. (2003) Interactions between tsetse and trypanosomes with implications for the control of trypanosomiasis. Adv Parasitol 53, 1–83

45. Roditi, I., and Lehane, M. J. (2008) Interactions between trypanosomes and tsetse flies. Curr Opin Microbiol 11, 345–351

46. Hirumi, H., and Hirumi, K. (1989) Continuous cultivation of *Trypanosoma brucei* blood stream forms in a medium containing a low concentration of serum protein without feeder cell layers. J.Parasitol. 75, 985–989

47. Hirumi, H., Doyle, J. J., and Hirumi, K. (1977) Cultivation of bloodstream *Trypanosoma brucei*. Bull World Health Organ 55, 405–409

48. Brun, R., and Shonenberger, M. (1979) Cultivation and in vitro cloning of procyclic culture forms of *Trypanosoma brucei* in a semi-defined medium. Acta Tropica 36, 289–292

49. Bolger, A. M., Lohse, M., and Usadel, B. (2014) Trimmomatic: a flexible trimmer for Illumina sequence data. Bioinformatics 30, 2114–2120

50. Liao, Y., Smyth, G. K., and Shi, W. (2014) featureCounts: an efficient general purpose program for assigning sequence reads to genomic features. Bioinformatics 30, 923–930

51. Robinson, M. D., McCarthy, D. J., and Smyth, G. K. (2010) edgeR: a Bioconductor package for differential expression analysis of digital gene expression data. Bioinformatics 26, 139–140

52. McCarthy, D. J., Chen, Y., and Smyth, G. K. (2012) Differential expression analysis of multifactor RNA-Seq experiments with respect to biological variation. Nucleic Acids Res 40, 4288–4297

53. Schmittgen, T. D., and Livak, K. J. (2008) Analyzing real-time PCR data by the comparative C(T) method. Nat Protoc 3, 1101–1108

54. Brenndorfer, M., and Boshart, M. (2010) Selection of reference genes for mRNA quantification in *Trypanosoma brucei*. Mol Biochem Parasitol 172, 52–55

55. Clemmens, C. S., Morris, M. T., Lyda, T. A., Acosta-Serrano, A., and Morris, J. C. (2009) *Trypanosoma brucei* AMP-activated kinase subunit homologs influence surface molecule expression. Exp Parasitol 123, 250–257

56. Field, M. C., Allen, C. L., Dhir, V., Goulding, D., Hall, B. S., Morgan, G. W., Veazey, P., and Engstler, M. (2004) New approaches to the microscopic imaging of *Trypanosoma brucei*. Microsc Microanal 10, 621–636

57. Shameer, S., Logan-Klumpler, F. J., Vinson, F., Cottret, L., Merlet, B., Achcar, F., Boshart, M., Berriman, M., Breitling, R., Bringaud, F., Butikofer, P., Cattanach, A. M., Bannerman-Chukualim, B., Creek, D. J., Crouch, K., de Koning, H. P., Denise, H., Ebikeme, C., Fairlamb, A. H., Ferguson, M. A., Ginger, M. L., Hertz-Fowler, C., Kerkhoven, E. J., Maser, P., Michels, P. A., Nayak, A., Nes, D. W., Nolan, D. P., Olsen, C., Silva-Franco, F., Smith, T. K., Taylor, M. C., Tielens, A. G., Urbaniak, M. D., van Hellemond, J. J., Vincent, I. M., Wilkinson, S. R., Wyllie, S., Opperdoes, F. R., Barrett, M. P., and Jourdan, F. (2015) TrypanoCyc: a community-led biochemical pathways database for *Trypanosoma brucei*. Nucleic Acids Res 43, D637–644

